# Functional analysis of the UVR8 photoreceptor from the monocot *Zea mays*

**DOI:** 10.1101/649905

**Authors:** Fernández María Belén, Lamattina Lorenzo, Cassia Raúl

## Abstract

Low UV-B fluence is a signaling stimulus that regulates various physiological processes and induces photomorphogenic responses in plants. The specific UV-B receptor UVR8 is a key component in these processes. Although UVR8 sequence is conserved, few homologs have been cloned and reported to be functional. Here we show the cloning and functional analysis of *Zea mays* UVR8 (*Zm*UVR8). *Zm*UVR8 presents 73% of identity with *At*UVR8, maintaining the key tryptophan responsible of UV-B perception. *Zm*UVR8 also contains the VP domain, involved in the interaction with the proteins CONSTITUTIVELY PHOTOMORPHOGENIC 1 (COP1) and REPRESSOR OF UV-B PHOTOMOPHOGENESIS 1 (RUP1). Whereas *UVR8* was expressed in non-irradiated *Arabidopsis* and maize leaves, after 2h of UV-B irradiation, its expression was reduced. The expression of chalcone synthase (*CHS*), involved in flavonoid biosynthesis and regulated by UVR8, was increased in irradiated *Arabidopsis* and maize leaves. *Arabidopsis uvr8-1* null mutant was complemented with *Zm*UVR8 driven by the CaMV-35S promoter and fused to eGFP. *Zm*UVR8-eGFP fusion was mainly localized in nuclei of transgenic lines, irrespective of UV-B treatments. UV-B suppressed hypocotyl elongation in WT *Arabidopsis* plants, whereas in *uvr8-1* hypocotyl growth was observed. However, hypocotyl elongation was reduced in UV-B irradiated transgenic lines complemented with *Zm*UVR8. Moreover, *CHS* and transcription factor *HY5* (ELONGATED HYPOCOTYL 5) expression were also restored in these plants. These results confirm that *Zm*UVR8 is similar enough to *At*UVR8 to restore UV-B perception and signaling in *Arabidopsis* mutant *uvr8-1*, thus being a functional UV-B photoreceptor.

## 1. Introduction

Ultraviolet-B (UV-B) radiation is the region between 280 and 315 nm of the total Sun’s electromagnetic spectrum. Ultraviolet irradiances reaching the surface of the Archean Earth were higher than the current ones because of the absence of a significant ozone atmosphere (Cnossen et al. 2007). UV-B is higher in terrestrial environments compared to the water column (Rozema et al. 2002). High UV-B doses damages DNA, proteins, lipids, cell membranes, photosynthetic machinery and induces the production of reactive oxygen species (ROS). This affects plant cell integrity and viability, leading to growth retardation and to a decrease in crop yield and quality (Brosché and Strid 2003; Frohnmeyer and Staiger 2003; Jordan 1996). Consequently, plants evolved mechanisms to avoid UV-B damage during the colonization of exposed habitats (Tilbrook et al. 2013). Ancient photosynthetic organisms like cyanobacteria and various eukaryotic algae, use mycosporine-like amino acids (MAAs) as protective compounds against UV-B (Llewellyn and Airs 2010; Rastogi and Incharoensakdi 2013; Rozema et al. 2002). Land plants co-evolved with environmental UV-B levels, and the complexity of UV-B absorbing molecules increased accordingly from algae to higher plants (Rozema et al. 2002). The protein UV RESISTANCE LOCUS (UVR8) was identified as the specific UV-B receptor in plants (Brown et al. 2005). UVR8 signaling significantly contributes to UV-B acclimation responses and the establishment of UV-B tolerance. UVR8 regulates metabolic and developmental processes and induces physiological and photomorphogenic responses. The usually UVR8-mediated UV-B responses are the inhibition of hypocotyl growth and the accumulation of flavonols and anthocyanins. However, nowadays it is suggested that additional physiological responses are modulated by UVR8: phototropism, thermomorphogenesis, circadian clock, auxin signaling, defense, salt stress tolerance, shade avoidance, chloroplast development, stomatal opening, leaf development and downward leaf curling (for a review see (Yin 2017)).

Chlorophytes are the earliest organisms with an UV-B photoreceptor, *Chlamydomonas reinhardtii* being the first organism to own a functional UVR8 (Fernandez et al. 2016; Han et al. 2019). Since its discovery in 2005 (Brown et al. 2005), cumulative evidence allows us to consider a landscape of UVR8 interactions. In the absence of UV-B, the UVR8 homodimer is mainly located in the cytoplasm. In the nucleus, the E3 Ubiquitin ligase COP1 (CONSTITUTIVELY PHOTOMORPHOGENIC 1) represses the activity of the transcription factor HY5 (ELONGATED HYPOCOTYL 5) (Favory et al. 2009). The transcription factor WRKY36 (WRKY DNA-BINDING PROTEIN 36) represses HY5 (Yang et al. 2018), and BES1 (BRI1-EMS-SUPPRESSOR1) and BIM1 (BES1-INTERACTING MYC-LIKE 1) induce BR-responsive gene expressions (Liang et al. 2018).

Following UV-B irradiation, UVR8 rapidly monomerizes and interacts in the nucleus with COP1 and WRKY36. This complex avoids the degradation of HY5, and induces the *HY5* expression. In turn, it triggers the UV-B regulated gene expression, leading to plant acclimation and stress tolerance (Heijde and Ulm 2012). Two of the HY5 upregulated genes are the Repressor of UV-B photomorphogenesis 1 and 2 (RUP1 and RUP2). Interaction of UVR8 with these proteins facilitates UVR8 dimerization, and subsequent inactivation (Heijde and Ulm 2012; Ulm and Jenkins 2015). UVR8 also interacts with BES1 and BIM1, inhibiting the brassinosteroid responsive genes, and reducing the hypocotyl elongation (Sun and Zhu 2018). Recently, it has been shown that the UVR8-COP1 complex participates in the PHYTCHROME INTERACTING FACTOR (PIF) 4 y 5 degradation, causing the UV-B-induced hypocotyl shortening (Sharma et al. 2019; Tavridou et al. 2019).

UVR8 is the first photoreceptor described that perceives UV-B through tryptophan residues instead of a prosthetic chromophore (O’Hara and Jenkins 2012; Ulm and Jenkins 2015).Two *Arabidopsis* UVR8 (*At*UVR8) high-resolution crystal structures have been determined using different crystallization conditions. However, they proved to be nearly identical in tertiary and quaternary structure (Christie et al. 2012; Wu et al. 2012; Yang et al. 2016; Zeng et al. 2015). *At*UVR8 contains a core domain that forms a seven-bladed β-propeller and a flexible C-terminal region of approximately 60 amino acids that contains a C27 region. Both the β-propeller domain and the C-terminal C27 domain of UVR8 are necessary and sufficient for interacting with COP1. Moreover, UVR8 interacts also with WRKY36 by its C-terminal (aminoacids 397 to 440) (Yang et al. 2018). *At*UVR8 has fourteen tryptophan residues, seven of which are exposed to the dimer interface. It also contains three conserved pentapeptide repeats with the motif “GWRHT” in blades 5, 6, and 7. This motif generates a closely clustered triad of tryptophans (W233, W285 and W337) which are the most important for UV-B photoreception (Christie et al. 2012; Wu et al. 2012; Zeng et al. 2015). Structural and mutagenesis studies show a primary role for W285 and W233in UV-B perception, whereas W337 is not essential in this process (Christie et al. 2012; Sun and Zhu 2018; Wu et al. 2012). The motif “GWRHT” from blade 6 contains W285 and is conserved in all UVR8 homologs analyzed (Fernandez et al. 2016; Han et al. 2019).

*Zea mays* also known as corn, is a cereal grain of agronomic importance, and has been used as a model organism in basic and applied research for nearly a century (Strable and Scanlon 2009). Previous work shows that increased UV-B radiation produces a significant reduction in dry matter accumulation and, consequently, affects yield. Moreover, an increase in flavonoid accumulation, a decrease in chlorophyll content in leaves and a reduction in protein level, sugar and starch of maize seeds have also been reported (Gao et al. 2004). In 2011, Casati et. al. described a transiently upregulation and subsequent downregulation of two UVR8-like genes upon UV-B exposure in maize leaves (Casati et al. 2011a; Casati et al. 2011b). However, these genes have little homology to *At*UVR8 as they were identified by homology to rice genome.

Although UVR8 is conserved, and sequences for this gene are found in all the *Viridiplantae* (Fernandez et al. 2016; Han et al. 2019), a few UVR8 homologs have been cloned and reported to be functional from green algae, moss and dicots. Up to now, there is no evidence of UVR8 photoreceptor from monocotyledon plants with confirmed functionality. Here, we report the molecular cloning, sequence and functional complementation of *Zm*UVR8, the UV-B receptor of *Zea Mays*.

## 1. Material and Methods

### 1.1. Plants material and growth conditions

Seeds of wild-type (WT) *Arabidopsis thaliana* ecotype *Landsberg erecta* (Ler) and the *uvr8-1* null mutant (Cloix et al. 2012) were kindly provided by Dr. Gareth Jenkins (University of Glasgow, Scotland). Seeds were surface sterilized in 30% (v/v) commercial bleach for one minute, rinsed with distilled sterile water and stratified for 72h at 4°C in darkness. Germinated seedlings were grown on agar plates containing half-strength Murashige and Skoog (MS) salts or in soil/ perlite/ vermiculite (3:1:1, v/v) under white light (160 μmol.m^-2^ s^-1^, fluorescent tubes) and long-day regime (light/dark: 16/8h) at 25°C in an environment controlled chamber.

Maize (*Zea mays* B73 inbreed line) seeds were kindly provided by Dr. Sofía Eugenia Olmos (INTA Pergamino, Argentina). Seeds were surface sterilized with 30% (v/v) commercial bleach for 20 minutes and rinsed in distilled water. Subsequently, seeds were germinated on water saturated filter paper in Petri dishes for 4-5 days and maintained at 25°C. Germinated seedlings were grown on soil/ vermiculite (3:1, v/v) under a long-day regime (light/dark: 16/8 h) at 25°C in an environment controlled chamber with white light at 160 μmol.m^-2^ s^-1^. The topmost leaf from V6 developmental stage (when sixth leaf are visible in the leaf whorl) plants was used for experiments.

### 1.2. Generation of transgenic *Arabidopsis* plants expressing *Zm*UVR8

The full length coding sequence from *Zm*UVR8 was amplified by PCR using the maize full-length EST ZM_BFb0066P22.r (Arizona Genomics Institute) as template, and the specific primers *Zm*UVR8-Fw and *Zm*UVR8Rv (Table 1). The amplified cDNA was cloned into the entry pENTR/D-TOPO vector and confirmed sequence, orientation and reading frame by DNA sequencing (Macrogen). The obtained entry clone was recombined with the Gateway pH7FWG2 binary destination vector for 35S-driven expression in plants, with C-terminal fusion to eGFP (35S::*Zm*UVR8-eGFP) (Karimi et al. 2002). This vector was introduced in the *Agrobacterium* strain GV3101 by electroporation (Koncz and Schell 1986). Transformation into *uvr8-1* mutant *Arabidopsis* was performed by floral dip (Clough and Bent 1998). Transformants were selected based on its ability to survive on half-strength Murashige and Skoog medium supplemented with 1% sucrose containing 15 mg L^-1^ hygromycin. Resistant seedlings were then transferred to soil and grown under conditions described above.

**Table 1.**
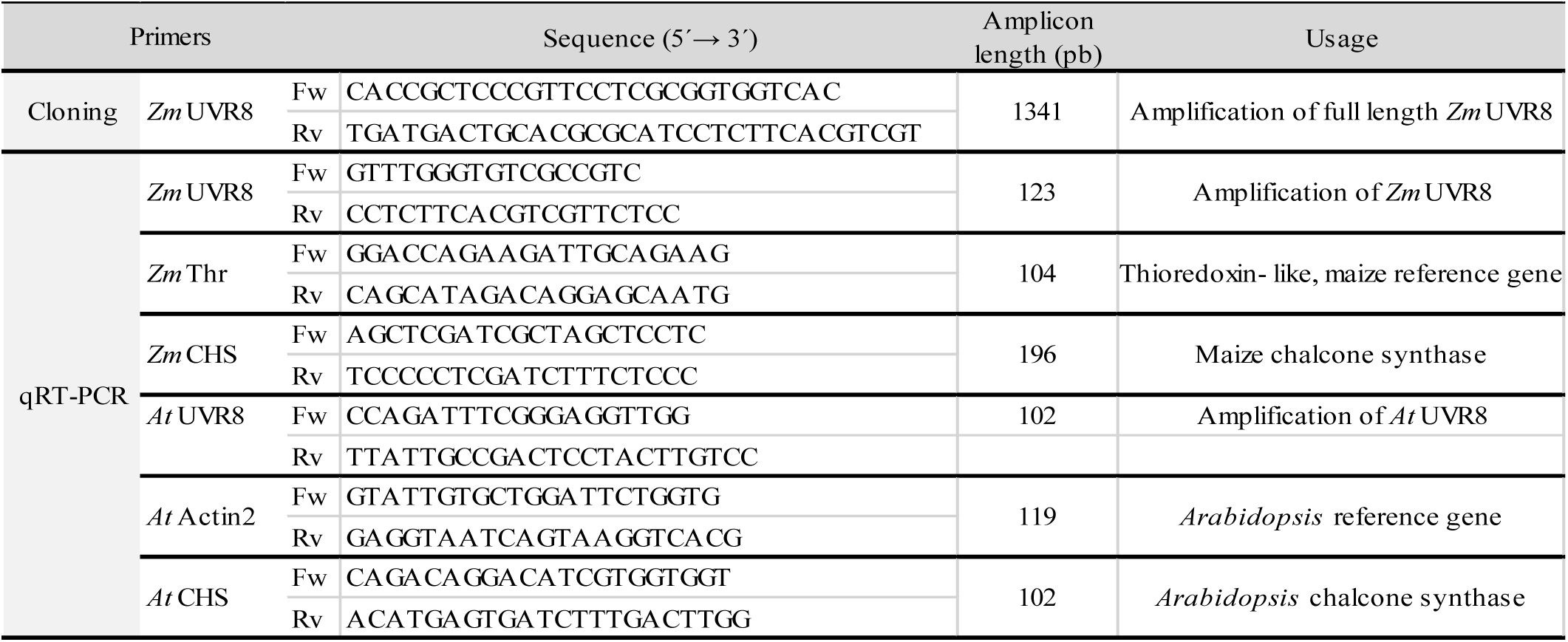
List of primers used in this work.

The transgenic lines generated were shown to have the transgene integrated at a single genetic locus through segregation analysis. Homozygous T3 (#6.5) and T4 (#5.1.7) independent lines were obtained by self-crossing and were used for the experiments. The level of transgene expression in each line was examined by qRT-PCR and immunoblot.

### 1.3. UV-B Treatments

For experiments involving UV-B light treatments, *Arabidopsis* seedlings were exposed 2h to white light (100 μmolm^-2^s^-1^, fluorescent tubes) supplemented with 3.47 μmol m^-2^s^-1^ narrowband UV-B (Philips TL 100W/01) in a controlled environment chamber. The UV-B dose used is similar to the radiation measured from sunlight at midday in Mar del Plata summer (38.0055° S, 57.5426° W).

Maize plants were exposed for 0, 2 and 4h to white light (100 μmol.m^-2^s^-1^, fluorescent tubes) supplemented with 8.81 μmol m^-2^s^-1^ narrowband UV-B (Philips TL 100W/01) in a controlled environment chamber.

The spectral irradiance was determined with an Ultraviolet-B photo-radiometer (Delta ohm HD2102.1).

### 2.4. Expression analysis

For gene expression quantification, plant materials were harvested, frozen in liquid nitrogen and grounded under RNase-free conditions. Total RNA was extracted using TRIzol method, and treated with DNase I (Invitrogen) at 37°C for 30 min, following the manufacturer’s instructions. Then, the RNA was reverse-transcribed using the M-MLV reverse transcriptase (Thermo) following the manufacturer’s instructions. cDNA obtained was used for quantitative RT-PCR using Power SYBR Green PCR mix and a StepOne machine (Applied Biosystems). Primers used are listed in Table 1. PCR conditions were: 10 min at 95 °C, 40 cycles of 15 s at 95 °C and 1 min at 60°C. After amplification, a melting curve analysis was performed, which resulted in a single product specific melting curve. Negative controls for cDNA synthesis and qRT-PCR reactions were included in all cases. LineReg program was employed for the analysis of gene expression (Ruijter et al. 2009). The transcript relative quantifications were determined from the ratio between the starting concentration value of analyzed mRNA and the reference genes actin2 for *Arabidopsis* samples or thioredoxin-like (Thr) for maize as previously reported (Casati and Walbot 2004). The data shown are representative of at least three independent experiments.

For protein assays, leaves were harvested into liquid nitrogen and proteins extracted in 100mM buffer KPO4 (pH 7.4), 1mM EDTA and a cocktail of protease inhibitors. The homogenate was centrifuged for 10 min a 10000 x g at 4°C. Protein concentration was determined by a Bradford assay. Thirty micrograms of total protein were loaded for *Zm*UVR8 protein expression and separated by 12% denaturing SDS-PAGE. Immunoblots were incubated with anti-GFP antiserum as primary antibody. After several washes, a secondary anti-mouse antibody conjugated to alkaline phosphatase and developed by NBT/BCIP staining. The membranes were stained with Ponceau S to reveal the Rubisco large subunit (rbcL) as loading control.

### 2.5. *Zm*UVR8- eGFP Subcellular Localization

Fifteen day old plants irradiated 1h with white light or white light supplemented with UV-B (3.47 μmol.m^-2^s^-1^), were vacuum infiltrated with 5 μg. mL^-1^ of Hoechst 33342 (Invitrogen Molecular Probes) in buffer PBS, 0.2% Triton-X100 for 4 min and maintained in shake at 50 rpm and darkness for 1h. Then, samples were washed 3 times with PBS. The subcellular localization of eGFP and Hoechst 33342 was visualized by a confocal laser scanning microscope (Nikon-C1siR Eclipse TiU) under oil (Biopack) with a ×40 objective. Images were taken using the Nikon EZ-C1 3.90 software. eGFP and Hoechst were excited using an argon laser at 488 nm and a laser at 408 nm, respectively. eGFP emission was collected between 515 and 530 nm to avoid crosstalk with chloroplast autofluorescence. Hoechst 33342 fluorescence was collected at 440/50 nm. The same microscope settings for GFP and Hoechst 33342 detection were used before and after UV-B illumination. Colocalization analysis was performed on two independent transgenic lines. The data shown are representative of at least three independent experiments.

### 2.6. Hypocotyl length measurement

Seedlings were grown for 5 days on agar plates of half strength Murashige and Skoog (MS) salts containing 1% sucrose in white light or white light supplemented with 3.47 μmol.m^-2^ s^-1^ UV-B. Photographs were taken after treatments and hypocotyl lengths were measured using the ImageJ software (http://rsb.info.nih.gov/ij). Three independent biological replicates were performed for all experiments using al least 10 seedlings for each replicate (n≥30).

### 2.7. Bioinformatic analysis

Multiple sequence alignments were performed using MAFFT server (http://mafft.cbrc.jp/alignment/server/) and edited with GeneDoc (Nicholas and Nicholas 1997).

## 1. Results

### 3.1. An *At*UVR8 homolog with a conserved structure is found in maize

UVR8 sequences from green algae (*C. reinhardtii* (Tilbrook et al. 2016)), moss *(P. patens* (Soriano et al. 2018)), liverworth (*M. polymorpha* (Soriano et al. 2018)) and dicots (*S. lycopersicum* (Li et al. 2018a), *M. domestica* (Zhao et al. 2016), *V. vinifera* (Liu et al. 2014), *B. platyphylla* (Li et al. 2018b) and *P. euphratica* (Mao et al. 2015)) have been cloned, characterized, and shown to restore the loss-of-function of the UVR8 null mutant *uvr8-1* (Kliebenstein et al. 2002)

However, no monocot functional UVR8 protein has been reported so far. To this end, the *Arabidopsis* protein sequence (AAD43920.1) was used as bait in a PSI-BLASTp for searching the UVR8 homolog in maize, using the NCBI (www.ncbi.nlm.nih.gov/) and Phytozome databases (www.phytozome.jgi.doe.gov/pz/portal.html). One sequence was found (GRMZM2G003565), herein named “*Zm*UVR8”. Figure 1A shows the comparison between *Zm*UVR8 and the other functional UVR8 homologs. *Zm*UVR8 has 443 amino acids length, a calculated molecular mass of 47.15 kDa, and 73% of identity to *At*UVR8. Key tryptophan residues responsible of UV-B perception (W233, 285 and 337) are conserved in *Zm*UVR8, as well as the VP domain in the C27 region, involved in the interaction with COP1 and RUP (Figure 1A). Figure 1B shows that *Zm*UVR8 has the same domain profile than *At*UVR8, including the conserved tryptophans, the C27 domain and the seven repeated RCC1 domains.

**Fig. 1.**
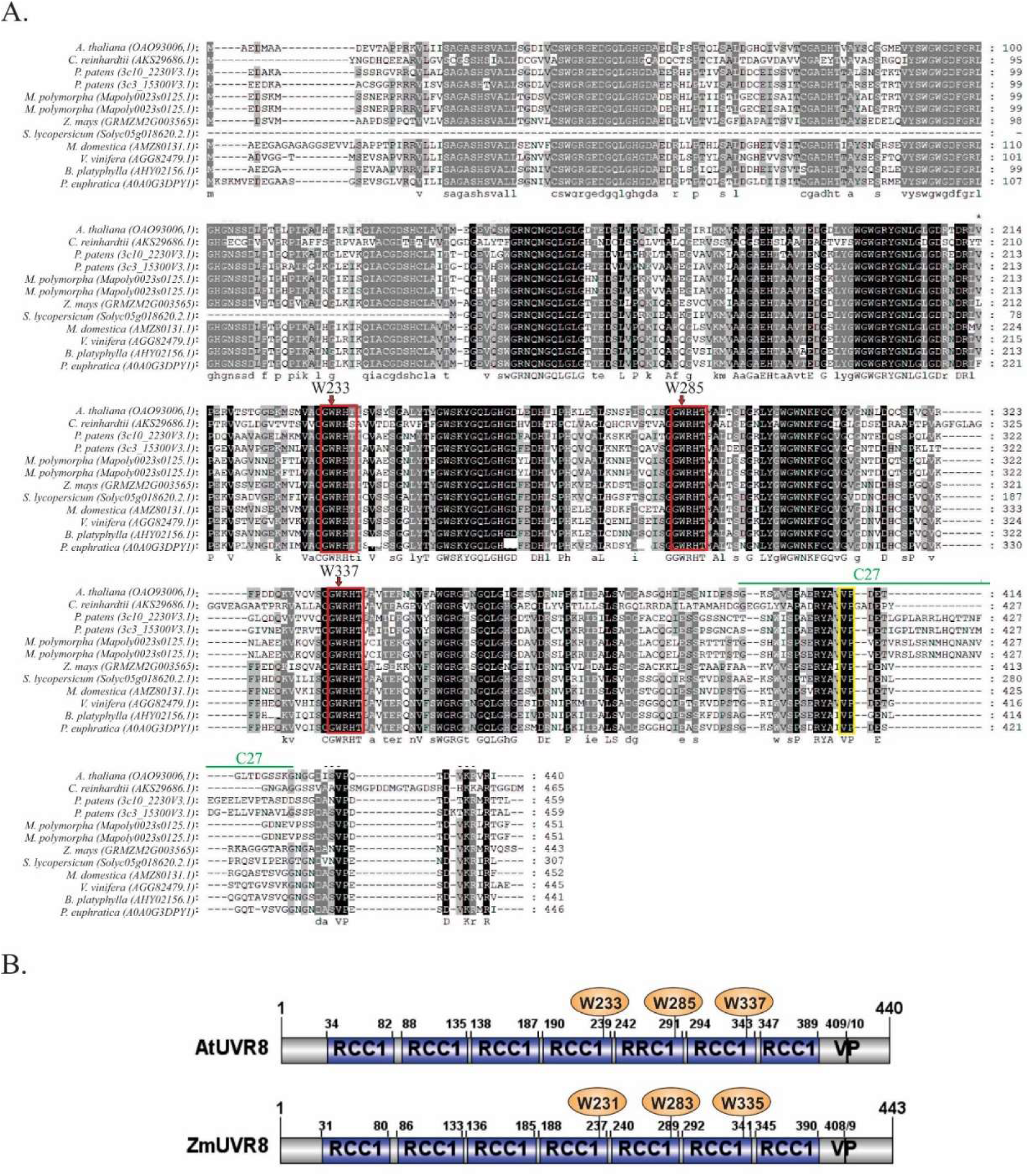
*Zm*UVR8 sequence and structural domains. **A.** Multiple sequence alignment of *At*UVR8 homologs. Sequences of proteins that restore the loss-of-function of the *uvr8-1* null mutant were aligned with MAFFT (http://mafft.cbrc.jp/alignment/server/) and edited with GeneDoc (Nicholas and Nicholas 1997). Red boxes indicate “GWRHT” motifs and the VP domain. The accession number of each sequence is given next to the species name. Conserved residues common to all sequences are shadowed in black and less identity is shown in gray scale. Capital letter indicates 100% of homology whereas lowercase indicates minor identity. The yellow box shows the “VP” domain in the C27 domain, important for interaction with COP1 and RUP proteins. **B)** Schematic representation of structural domains of *At*UVR8 and *Zm*UVR8 proteins. Analysis of amino-acid sequences was performed at the National Center for Biotechnology Information (NCBI) database. RCC1 (pfam00415), Regulator of chromosome condensation (RCC1) repeat. Arrows indicate significant tryptophans in UV-B perception (W233, 285 and 337). VP: Valine-Proline domain.

Blastp analysis in Table 2 shows that sequences with high percentage of identity with the components of the *Arabidopsis* UVR8 signaling pathway were found in maize: COP1 (70%), HY5 (68%), HYH (48%), RUP1 (45%), RUP2 (46%), WRKY36 (35%), BES1 (50%) and BIM1 (41%). These results indicate that *Zm*UVR8 could be a functional UV-B receptor, triggering a UVR8 signaling cascade in maize.

**Table 2.**
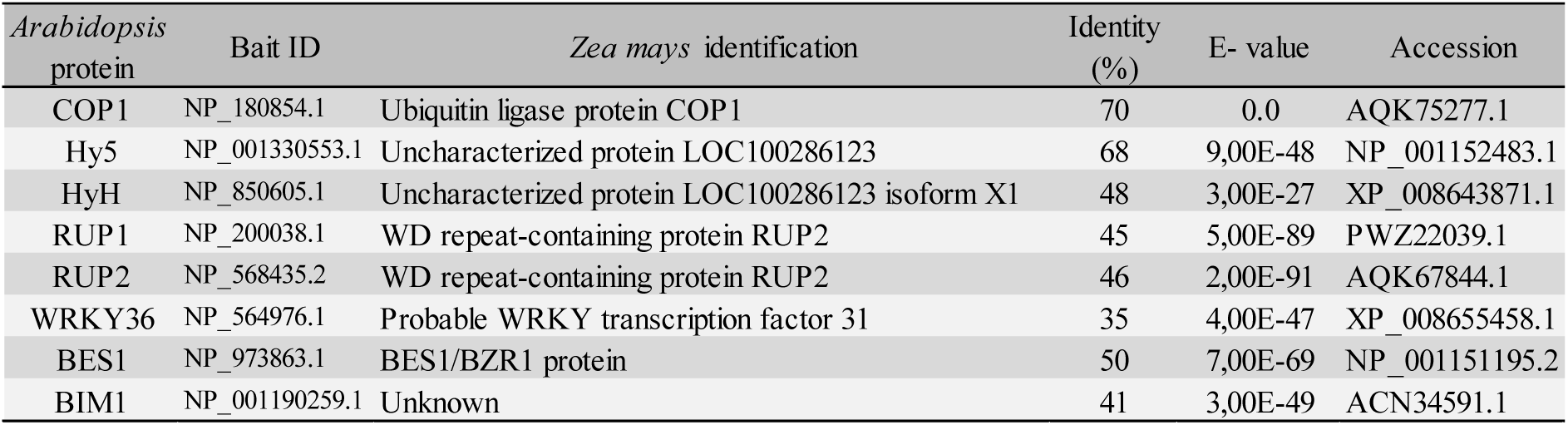
Identification of maize homologs to the *Arabidopsis* UVR8 signaling pathway. Proteins involved in UVR8 signaling cascade from *Arabidopsis* were used as bait in BLASTp analysis restricting the search to *Z. mays* and the non-redundant protein sequences database from NCBI.

### 3.2. *Zm*UVR8 and *At*UVR8 expression are downregulated by UV-B irradiation

Previous studies show that *At*UVR8 is a ubiquitously expressed gene, allowing an immediate reaction of plants to UV-B exposure (Favory et al. 2009; Rizzini et al. 2011). UVR8 expression was analyzed in *Arabidopsis* and maize plants irradiated with 3.47 and 8.81 μmol.m^-2^ s^-1^ of UV-B respectively. Figure 2 shows quantitative RT-PCR analysis of UVR8 expression in irradiated and non-irradiated leaves. After 2h of UV-B irradiation, UVR8 expression was reduced 3-fold in *Arabidopsis* and 6-fold in maize, respectively. However, CHS expression was increased in both species after UV-B. These results indicate a similar regulation of UVR8 and CHS expression under UV-B treatment both in *Arabidopsis* and maize plants.

**Fig. 2.**
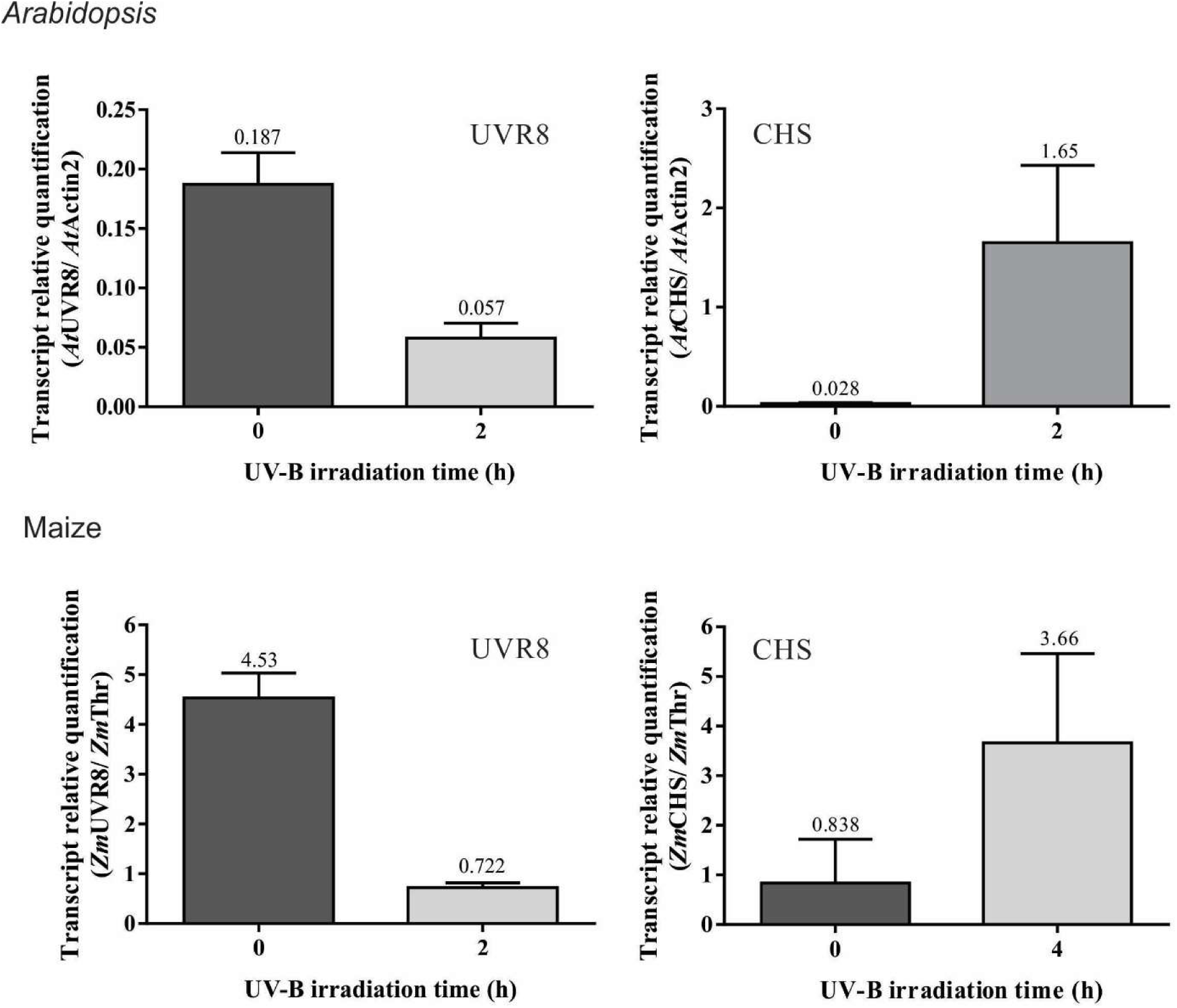
Expression of UVR8 and CHS in response to UV-B in *Arabidopsis* and maize plants. UVR8 and CHS transcript levels were analyzed by qRT-PCR. Three week–old *Arabidopsis* and maize plants were irradiated during 2 and 4h with 3.47 and 8.81 μmol.m^-2^ s^-1^ of UV-B respectively. Non-irradiated plants were used as control. Expression of Actin2 and thioredoxin-like (Thr) were used for *Arabidopsis* and maize normalization respectively. Primers are listed in Table 1. Error bars indicate the standard deviation over three biological replicates (n=3).

Table 3 compares the expression of *ZmUVR8* with those of the other functional homologues. Whereas UV-B has no effect on *UVR8* expression in *M. polymorpha* and *V. vinifera* grapes, it upregulates transcript levels in different tissues from *M. domestica* and *B. platyphylla*. These findings, together with results shown in Figure 2, where *At*UVR8 and *Zm*UVR8 transcript levels were downregulated by this radiation, suggest the existence of different regulatory mechanisms for *UVR8* expression in different plant species and tissues.

**Table 3.** *UVR8* expression after UV-B stress in plant with confirmed UVR8 functional homologs.

| Specie | Tissue | UVR8 expression in respopnse to UV-B | UV-B dose | Reference |
| --- | --- | --- | --- | --- |
| *Arabidopsis thaliana* | Leaf | Downregulated after 2h of treatment | 3.47 $\mu\text{mol m}^{-2} \text{s}^{-1}$ | Figure 2 |
| *Chlamydomonas reinhardtii* | nd |  |  | Tilbrook et. al., 2016 |
| *Physcomitrella patens* | nd |  |  | Soriano et. al., 2018 |
| *Marchantia polymorpha* | Plant | No significant effect of treatment after 3 | 3 $\mu\text{mol m}^{-2} \text{s}^{-1}$ | Soriano et. al., 2018<br>Kondou et. al., 2019 |
| *Zea mays* | Leaf | Downregulated after 2h of treatment | 8.81 $\mu\text{mol m}^{-2} \text{s}^{-1}$ | Figure 2 |
| *Solanum lycopersicum* | nd |  |  | Li et. al., 2018 |
| *Malus domestica* | Fruit skin | Upregulation after 6h of treatment | 1.5 $\mu\text{mol m}^{-2} \text{s}^{-1}$ | Zhao et. al, 2016 |
| | Apple callus | Upregulation after 1h of treatment | 1.5 $\mu\text{mol m}^{-2} \text{s}^{-1}$ | |
| *Vitis vinifera* | Fruit | Not influenced by UV-B | Field radiation | Liu et. al., 2014 |
| *Betula platyphylla* | Tissue culture seedling | Upregulation after 6h of treatment | 1.5 $\mu\text{mol m}^{-2} \text{s}^{-1}$ | Li et. al., 2018 |
| *Populus euphratica* | nd |  |  | Mao et. al., 2015 |

Nd: not determined.

### 3.3 Transgenic *Zm*UVR8 expression in the *uvr8-1* null mutant

For insights into the *in vivo* role of *ZmUVR8*, we cloned the *ZmUVR8* cDNA into the pH7FWG2 plant expression vector driven by the CaMV-35S promoter and fused to eGFP (35S::*Zm*UVR8-eGFP). This construct was used to transform the *Arabidopsis uvr8-1* null mutant. After repeated selection on hygromycin and microscopy eGFP-screening, two independent T3 and T4 homozygous lines (#6.5 and #5.1.7 respectively) were obtained. *ZmUVR8* expression was determined by quantitative RT-PCR using specific primers that did not amplify *At*UVR8 (Online resource S1). Figure 3A illustrates that both lines expressed *ZmUVR8* mRNA, line #6.5 reaching the highest level. Immunoblot with anti-GFP antibody in Figure 3B shows that *Zm*UVR8-eGFP was present in both transgenic lines.

**Fig. 3.**
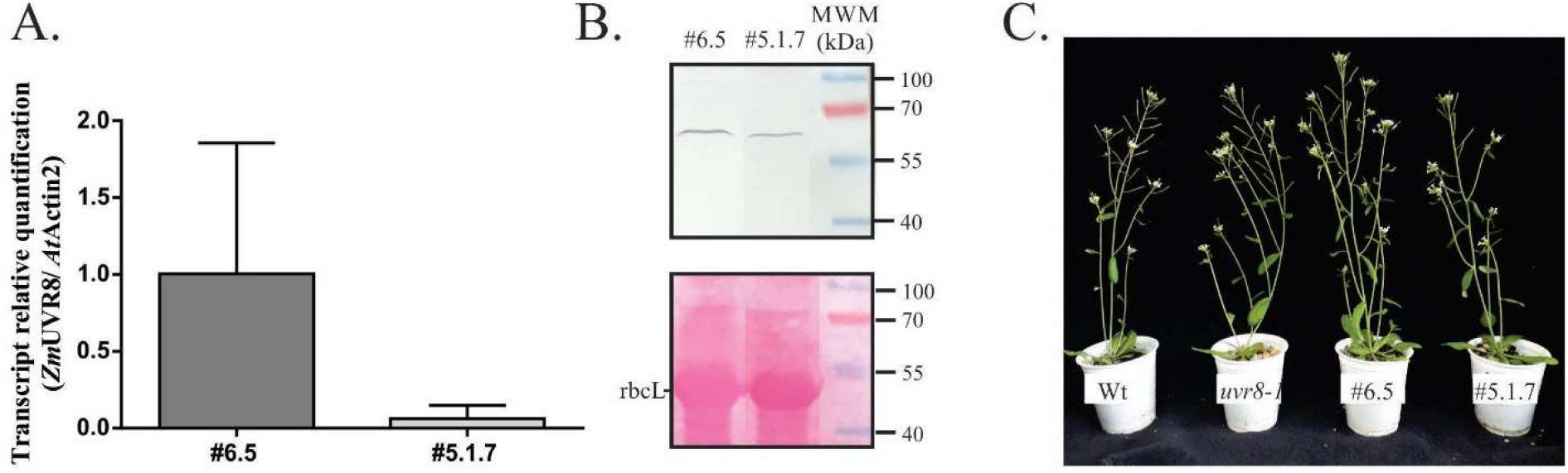
35S::*Zm*UVR8-eGFP expression analysis in two *Arabidopsis* homozygous lines. **A.** Leaves from three-weeks-old transgenic *Arabidopsis* plants were used to analyze *Zm*UVR8 transgene transcript level by qRT-PCR. Expression of Actin2 was used for normalization. Primers are listed in Table 1. Error bars indicate the standard deviation over three biological replicates. **B.** Denaturating immunoblot from *Arabidopsis* transgenic lines #6.5 and #5.1.7 protein extracts (upper panel). Immunodetection was performed using anti-GFP antiserum. Stained Rubisco large subunit (rbcL) is shown as a loading control (lower panel). **C.** Thirty-days-old WT, *uvr8-1* mutant and #6.5 and #5.1.7 transgenic *Arabidopsis* lines.

Online resource S2 shows no positive GFP signal in WT lines, ensuring the specificity of *Zm*UVR8-eGFP detection. Figure 3C shows that *Zm*UVR8 has no detrimental effects on plant development, because no evident phenotypic differences were found among WT, *uvr8-1*, #6.5 and #5.1.7, after 30 days.

The sub-cellular localization of *Zm*UVR8-eGFP, was analyzed using a confocal laser scanning microscope. Hoechst 33342 staining was employed for nuclear staining. Figure <u>4</u> shows that *Zm*UVR8-eGFP fusion was mainly localized in nuclei of #6.5 and #5.1.7 lines, irrespective of UV-B treatments.

**Fig. 4.**
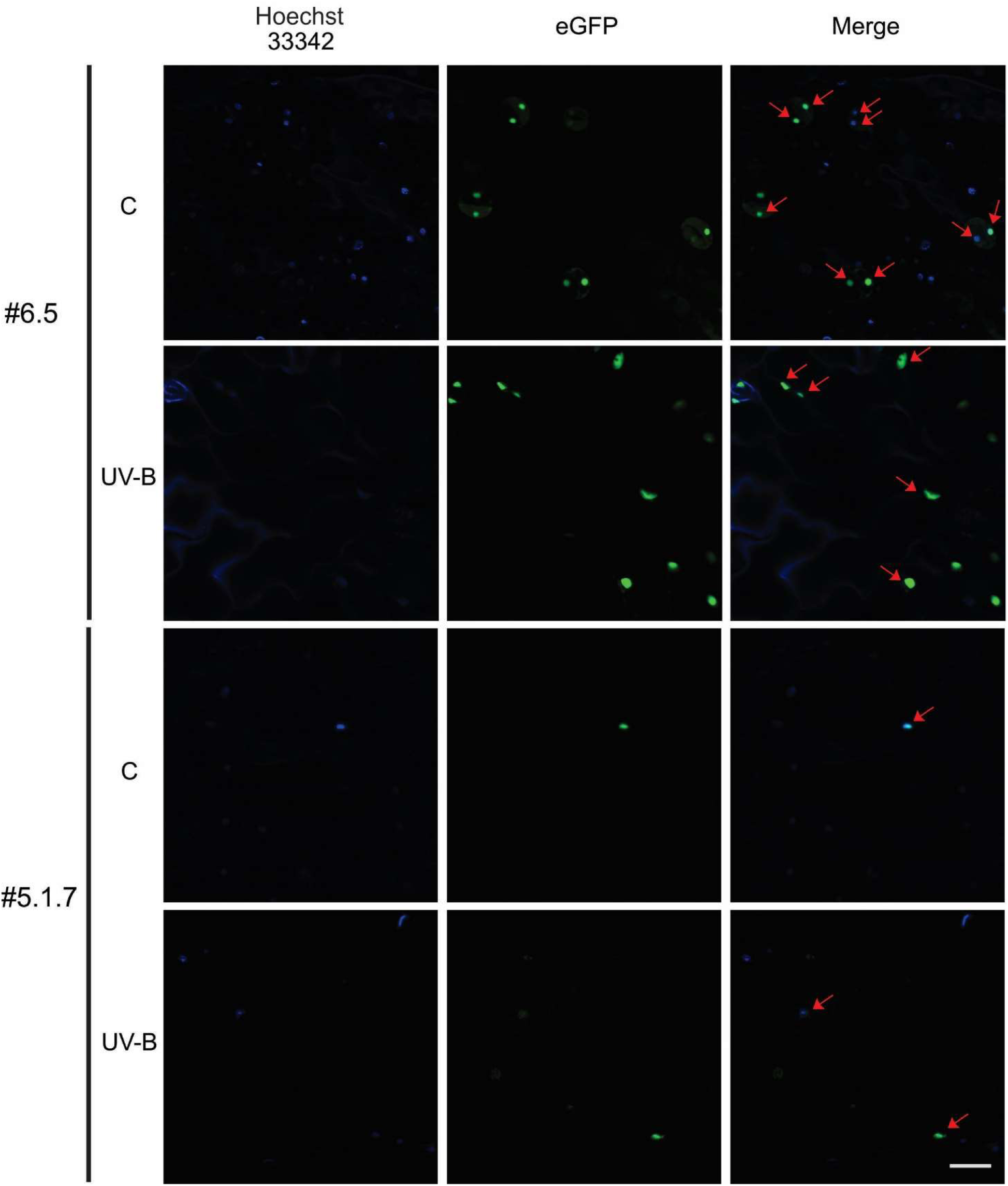
Subcellular localization of eGFP-*Zm*UVR8. Cellular localization of *Zm*UVR8-eGFP fusions revealed by confocal laser microscopy. Two-weeks-old *Arabidopsis* plants were irradiated with 100 μmolm^-2^s^-1^ of white light (C), or white light plus 3.47 μmol.m^-2^s^-1^ UV-B (UV-B) for 1 h. Hoechst 33342 was vacuum infiltrated to specifically stain nuclei. eGFP and Hoechst were excited using a laser at 488 nm and 408 nm, respectively. eGFP emission was collected between 515 and 530 nm to avoid crosstalk with chloroplast autofluorescence. Hoechst 33342 fluorescence was collected at 440/50 nm. The same microscope settings for GFP and Hoechst 33342 detection were used before and after UV-B illumination. Bar: 25 µm. #6.5 and #5.1.7: transgenic *Arabidopsis* lines expressing *Zm*UVR8-eGFP. Arrows indicate co-localization.

### 3.4. *Zm*UVR8 complements the *Arabidopsis uvr8-1* null mutant

The hypocotyl length inhibition is frequently used to analyze photoreceptors functionality in *Arabidopsis*. UV-B suppresses hypocotyl growth in wild-type *Arabidopsis* plants, whereas the *uvr8-1* mutant is impaired in this response (Favory et al. 2009). **E**xpression of WT *At*UVR8 fused to GFP or yellow fluorescent protein (YFP) in *uvr8-1* plants, restores the WT phenotype (Favory et al. 2009; Huang et al. 2014; O’Hara and Jenkins 2012). The recovering of this morphogenic response was established as a parameter of functionality for several UVR8 homologs (Kondou et al. 2019; Li et al. 2018a; Li et al. 2018b; Liu et al. 2014; Mao et al. 2015; Soriano et al. 2018; Tilbrook et al. 2016; Zhao et al. 2016). Figure 5 shows no differences among hypocotyl length of WT, *uvr8-1* and transgenic lines grown under white light. Although #5.1.7 line grown under white light appears slightly shorter than *uvr8-1* and #6.5 at the seedling stage, no differences were observed in 30-days-old plants (Fig. 3C). Figure 5 B shows that, under UV-B irradiation, the reduction in hypocotyl length was of 54% in WT plants. In contrast, the reduction in *uvr8-1* mutant was only of 31%. However, hypocotyl elongation was reduced 50% in line #6.5 and 60% in line #5.1.7, similar to WT plants (Fig. 5B). These results indicate that *Zm*UVR8 restore the *uvr8-1* null mutant phenotype under UV-B radiation demonstrating that is a positive regulator in UV-B induced photomorphogenesis.

**Fig. 5.**
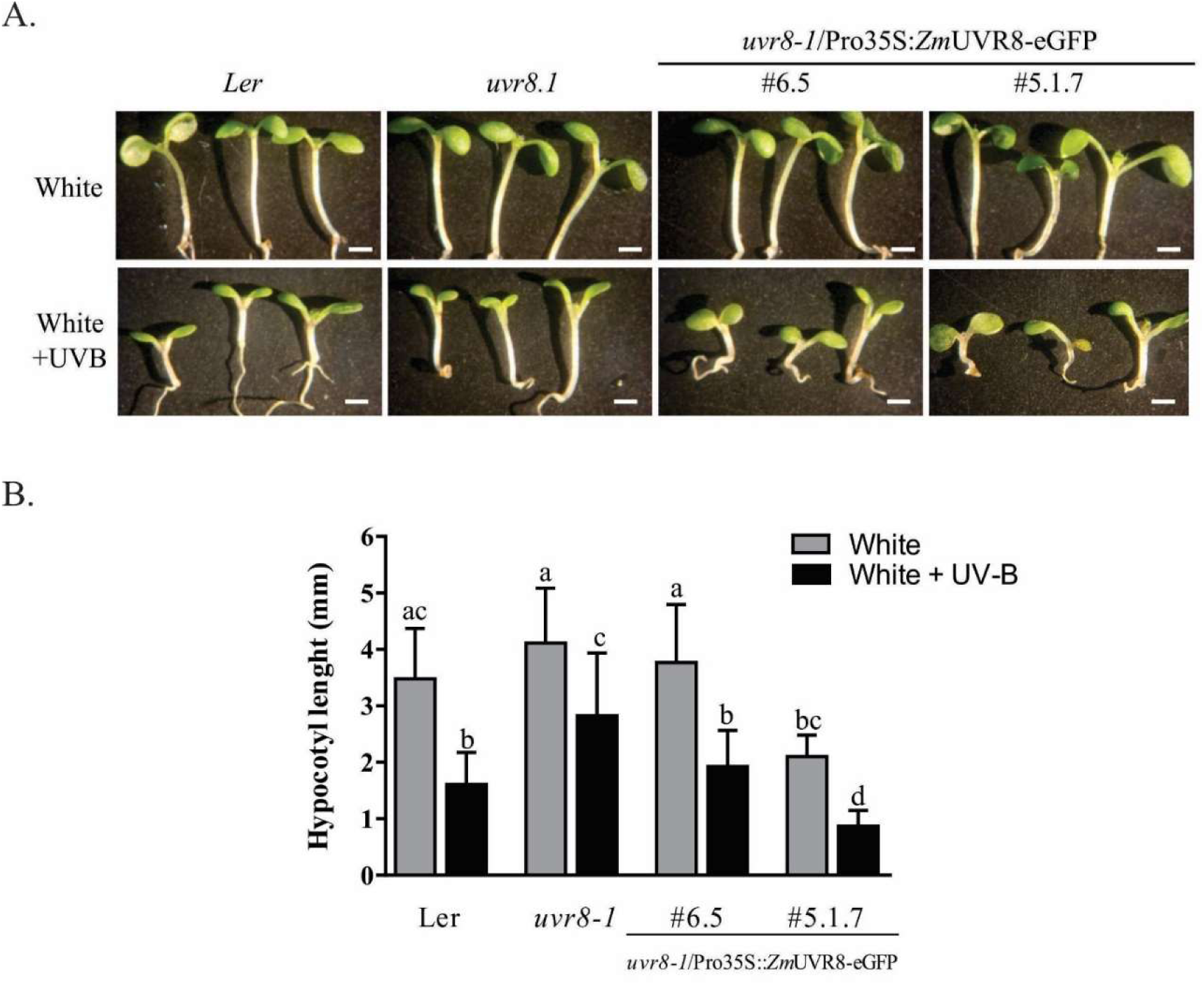
Functional complementation assay of eGFP-*Zm*UVR8 expressed in *Arabidopsis uvr8-1* mutant. **A.** Phenotypes of the *Arabidopsis* WT, *uvr8-1*, and #6.5 and #5.1.7 lines grown under 100 μmolm^-2^s^-1^ of white light, or white light plus 3.47 μmol.m^-2^ s^-1^ of UV-B. The scale bar represents 1mm. Photographs were taken after treatments. **B.** The hypocotyl lengths were measured using the ImageJ software (http://rsb.info.nih.gov/ij). Grey bars: white light, black bars: white light plus UV-B. Error bars indicate standard deviation. Different letters show significant differences between treatments (Kruskal-Wallis One-way ANOVA on ranks. Multiple comparisons: Dunn’s method, p<0.05). Three independent biological replicates were performed, n≥30.

### 3.5. *Zm*UVR8 promotes CHS and HY5 gene expression in *uvr8-1 Arabidopsis* mutant

In response to UV-B, plants increase the concentration of UV-B-absorbing phenolic compounds such as flavonoids. The enzyme chalcone synthase (CHS) catalyzes the first step of flavonoid biosynthesis and is regulated by the UVR8 signaling pathway (Jenkins 2014; Yonekura-Sakakibara et al. 2019). UVR8-COP1-SPA complexes induce the expression of the transcription factor HY5, which is a positive regulator of *CHS* expression (Jenkins 2014). In order to determine if *Zm*UVR8 is able to induce *CHS* and *HY5* expression in response to UV-B, three- weeks- old Ler, *uvr8-1* and #6.5 transgenic line plants were irradiated with white light or white light supplemented with UV-B for 2 hours. *CHS* and *HY5* expression was determined by quantitative RT-PCR. Figure 6 illustrates that, under white light irradiation, *CHS* and *HY5* have low basal expression in all plants. After UV-B irradiation, the expression of both genes was increased in WT plants, but the *uvr8-1* mutant was impaired in this response. However, UV-B-irradiated #6.5 line increased *CHS* and *HY5* expression (Fig. 6). These results, demonstrate that *Zm*UVR8 was able to trigger the UVR8 signaling in response to UV-B.

**Fig. 6.**
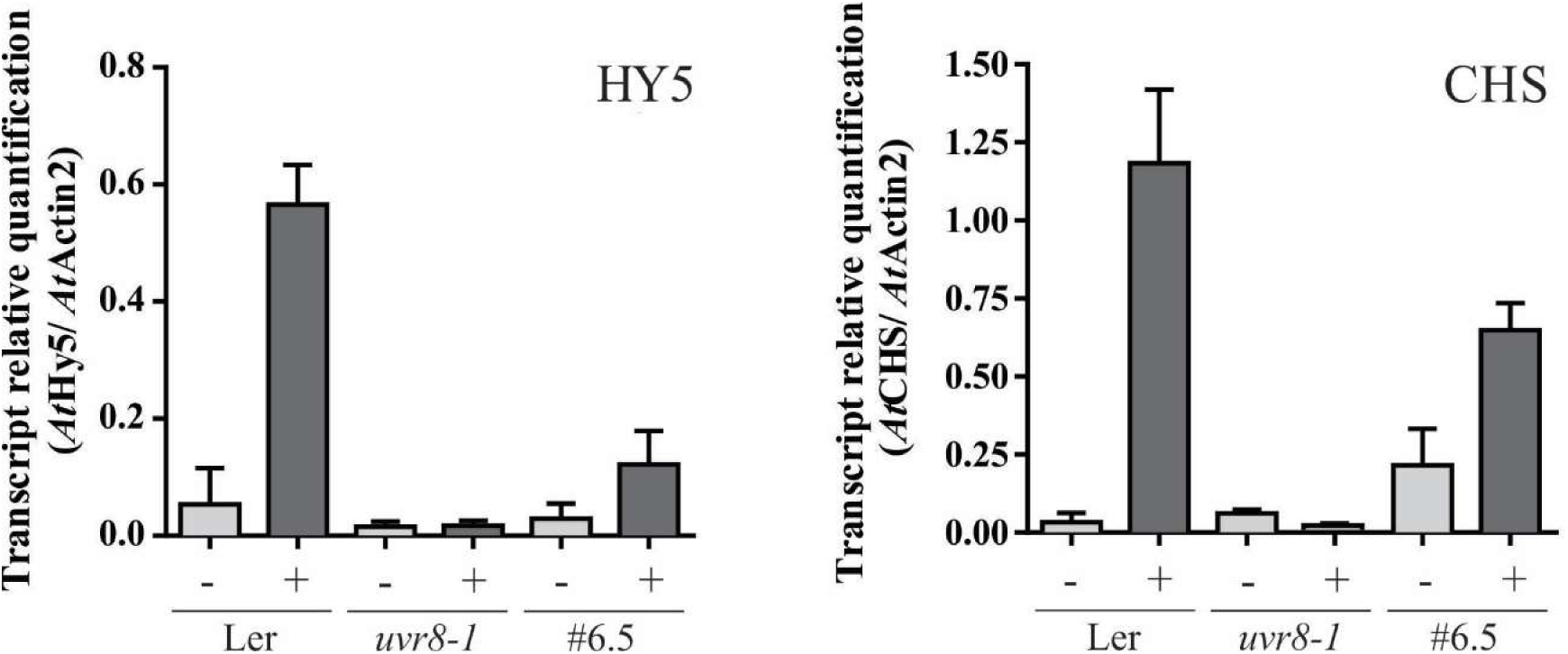
Expression levels of *CHS* and *HY5* in response to UV-B in WT, *uvr8-1* mutant and transgenic line #6.5 under UV-B irradiation. Three-weeks-old WT, *uvr8-1* and #6.5 transgenic line plants were exposed to white light, or white light supplemented with 3.47 μmol.m^-2^ s^-1^ of UV-B for 2h. The expression of *CHS* and *HY5* in leaves was determined by qRT-PCR. Expression of Actin2 was used for normalization. Error bars indicate the standard deviation of at least two biological replicates.

## Discussion

In this work we have cloned the maize *Zm*UVR8, the first functional UVR8 protein from monocots homologue to the *Arabidopsis* UV-B receptor.

Two maize UV-B-responsive genes were previously reported as UVR8-like homologs by comparison with rice genome (Casati et al. 2011a; Casati et al. 2011b). However, these genes present less than 34% of identity with *At*UVR8, and tryptophans 233 and 285, involved in UV-B perception, are not conserved (not shown). After the maize genome sequencing was completed, we found *Zm*UVR8 and, by phylogenetic studies, we concluded that a unique UVR8 sequence was present in maize (Fernandez et al. 2016). *Zm*UVR8 has a conserved sequence and a modular structure, in concordance with several UVR8 homologs (Fernandez et al. 2016). In addition, we found that maize proteins involved in UVR8 signaling have more than 35% of identity with those from *Arabidopsis* (Table 2). Considering that 30% of identity is enough to infer that two proteins are homologous (Pearson 2014), we suggest that the UVR8 signaling cascade might be present in maize. Furthermore, phylogenetic studies demonstrate that UVR8, SPAs and HY5 proteins are conserved from chlorophytes to angiosperms (Fernandez et al. 2016; Han et al. 2019). Functional studies will be necessary to confirm these hypotheses.

Although *UVR8* was reported as constitutive in *Arabidospsis*, we show herein that *UVR8* transcript decreased both in *Arabidopsis* and maize leaves after 2 hours of UV-B radiation. Likewise, *Betula platyphylla* UVR8 (*BpUVR8*) expression is induced by UV-B, and decreases after 9h of continuous irradiation (Li et al. 2018b). On the other hand, *UVR8* expression is increased by UV-B in the hypocotyls of radish sprouts (Wu et al. 2016). Variations in *UVR8* expression were also reported in UV-B-treated fruit skin and apple callus (Mao et al. 2015). Differences observed in *UVR8* expression in the literature could result from the different UV-B intensities used in the experiments and tissues analyzed. *At*UVR8 is a stable protein (Heilmann and Jenkins 2013) and a huge amount of UVR8 may influence the activity of COP1 and RUP as E3 ligases (Ren et al. 2019). It may be possible that after UVR8 was translated, gene expression could be decreased to avoid UVR8 overaccumulation in *Arabidopsis* and maize.

Little is known about the existence of different regulatory mechanisms for *UVR8* expression. Wu et. al. (2016) proposed that UV-B enhances the production of H2O2, thus increasing the level of NO to further magnify the *UVR8* expression. It was recently reported that *At*UVR8 expression may be modulated by blue light in UV-B-irradiated *Arabidopsis*. Cryptochrome 1 (Cry1) mutant *hy4* shows reduced *UVR8* expression in response to UV-B, suggesting a linkage between cry1, *UVR8* and UV-B (Khudyakova et al. 2019). Besides, it has been observed that *ZmUVR8* expression is increased by waterlogged in non-UV-B-irradiated maize root (Rajhi et al. 2011).

We obtained two independent homozygous *Arabidopsis* lines (#6.5 and #5.1.7), where *Zm*UVR8- eGFP complemented the *uvr8-1* null mutant both in hypocotyl elongation assay and induction of UVR8-regulated genes. *Arabidopsis* GFP-UVR8 shows increased nuclear localization following UV-B treatment of plants (Kaiserli and Jenkins 2007), but *Zm*UVR8-eGFP was constitutively located in nuclei in #6.5 and #5.1.7 lines.

No obvious nuclear localization signal (NLS) is found in UVR8, and there is no consensus about the mechanism of UVR8 translocation (Yin 2017). Yin et. al. (2016) reported that UVR8 interaction with COP1 is necessary for UVR8 translocation to nucleus, as well as for UVR8 signaling in response to UV-B (Yin et al. 2016). Moreover, it was demonstrated that the presence of UVR8 in the nucleus is necessary but not sufficient for its function. Kaiserli & Jenkins (Kaiserli and Jenkins 2007) fused a NLS to GFP-UVR8 and observed that the constitutive nuclear localization of NLS-GFP-UVR8 is insufficient to promote *HY5* expression in the absence of UV-B. Therefore, even though *Zm*UVR8 has nuclear localization irrespective of UV-B treatment, this radiation is necessary for downstream signaling as previously reported for *At*UVR8.

UV-B irradiation is not a mere stress signal but can also serve as an environmental stimulus to direct growth and development. A well-established UV-B morphogenic effect is the reduction of hypocotyl elongation (Kim et al. 1998). The hypocotyl growth of the *uvr8-1* mutant seedlings, in stark contrast to wild-type seedlings, was not inhibited by UV-B (Favory et al. 2009). The restoring of this morphogenic response was established as a parameter of complementation by functional UVR8 homologs (Kondou et al. 2019; Li et al. 2018a; Li et al. 2018b; Liu et al. 2014; Mao et al. 2015; Soriano et al. 2018; Tilbrook et al. 2016; Zhao et al. 2016).

*Zm*UVR8 restored the impaired UV-B hypocotyl growth suppression, and the *CHS* and *HY5* expression in *uvr8-1*, working as an effective component of the UVR8 pathway. These results demonstrate that similar signaling responses to UV-B are present in monocots and dicots plants. Consequently, a canonical UVR8 pathway could be present in monocotyledons.

## Conclusion

We identified the *ZmUVR8* gene from *Zea mays* (*Zm*UVR8) using multiple sequence comparison analysis. *ZmUVR8* gene expression is modulated by UV-B. The *Zm*UVR8 protein is highly conserved and has proven to be a functional homolog of the *Arabidopsis* UV-B receptor *At*UVR8.

## Acknowledgements

We acknowledge Dr. Rius from CEFOBI institute for technical support with maize samples.

## Supplementary figures

**Fig. S1.**
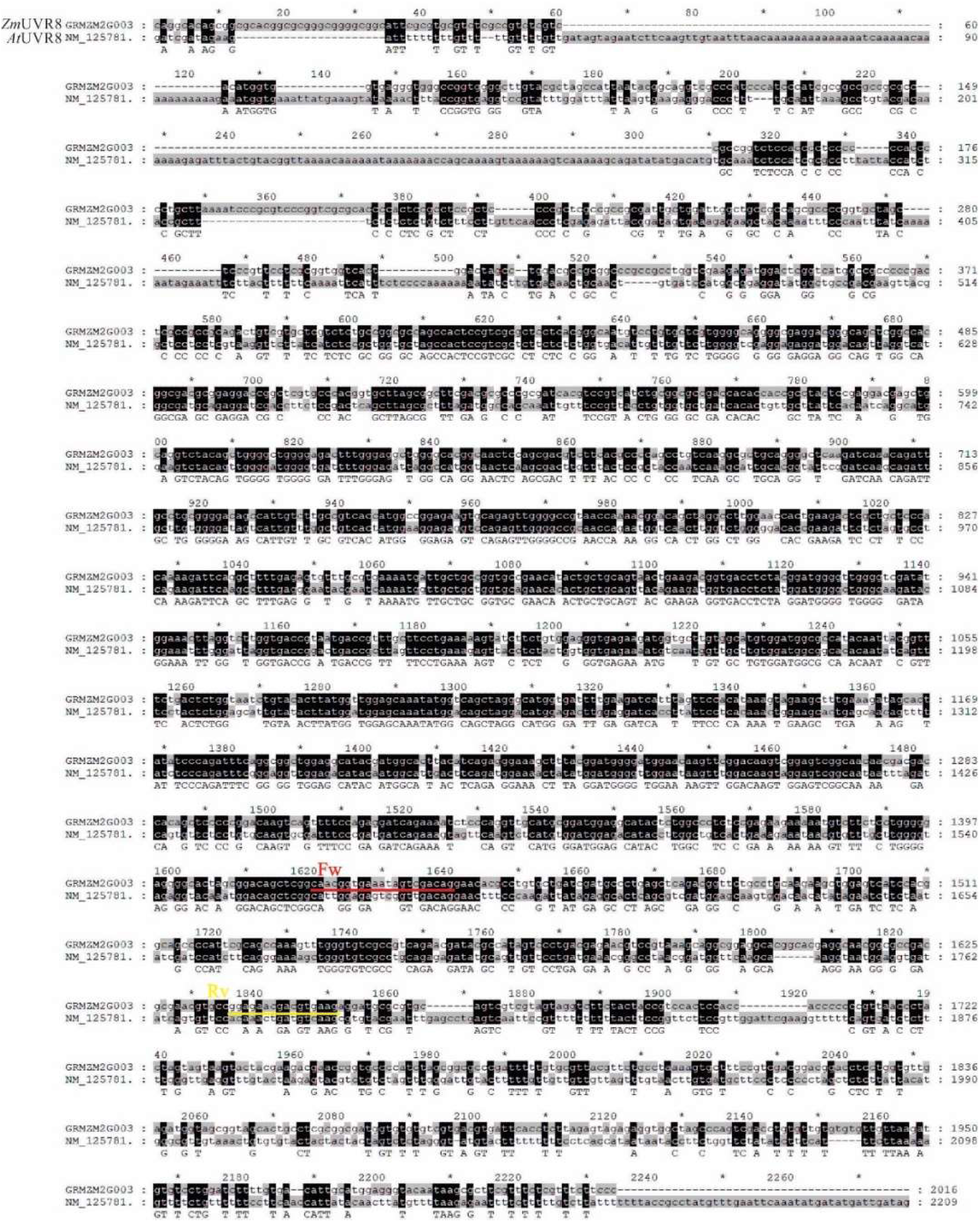
Sequence alignment of *At*UVR8 and *Zm*UVR8 transcripts. UVR8 transcript sequences from maize (GRMZMG003565_T03) and *Arabidopsis* (NM_125781.4) were aligned using MAFFT software (http://mafft.cbrc.jp/alignment/server/) and edited with GeneDoc (Nicholas and Nicholas 1997). Forward (Fw) and reverse (Rv) *Zm*UVR8 primers used for quantitative RT-PCR are shown in red and yellow respectively. Conserved residues common to both sequences are shadowed in black and less identity is shown in gray scale. Capital letter indicates 100% of homology whereas lowercase indicates minor identity.

**Fig. S2.**
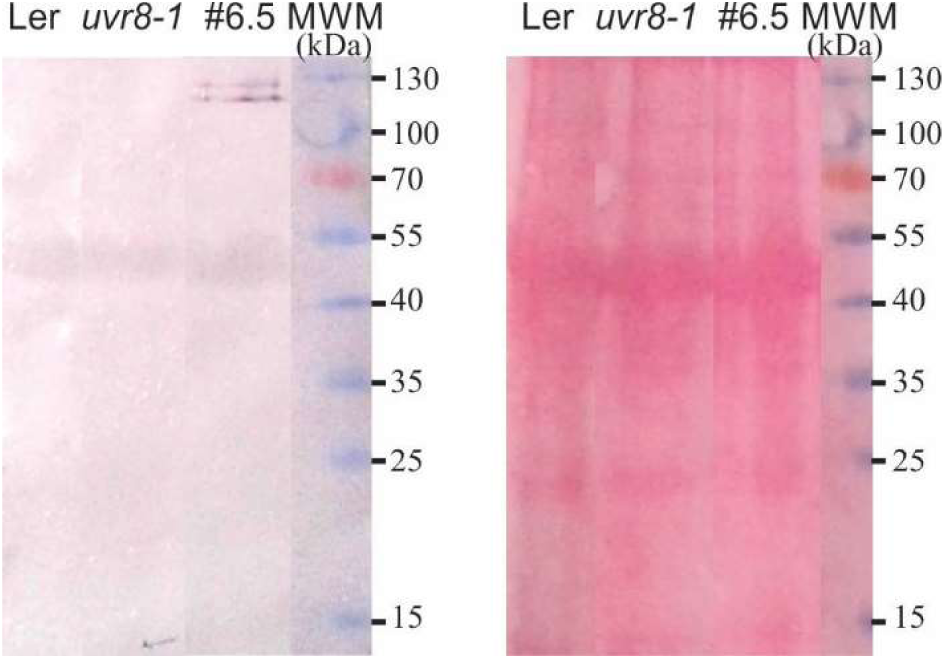
*Zm*UVR8 detection by immunoblot. Non-boiled total protein extracts from Ler, *uvr8-1* and #6.5 transgenic line were separated on 12%SDS-PAGE and immunoblot was probed with anti-GFP antibody (left panel). Ponceau staining is shown as loading control (right panel). MWM: molecular weight markers.

## Author Contributions

MF conducted experiments, interpreted data, drew figures, and collaborated in writing the manuscript. RC conceived the project and wrote the paper. LL supervised and improved the manuscript.

## Notes

#### Summary of Updates

This version was controlled with anti plagarism softwares

